# *Arap1* Loss Causes RPE Phagocytic Dysfunction and Subsequent Photoreceptor Death

**DOI:** 10.1101/2021.08.09.455745

**Authors:** Andy Shao, Antonio Jacobo Lopez, JiaJia Chen, Addy Tham, Seanne Javier, Alejandra Quiroz, Sonia Frick, Edward M. Levine, K.C. Kent Lloyd, Brian C. Leonard, Christopher J. Murphy, Thomas M. Glaser, Ala Moshiri

## Abstract

**Purpose:** *Arap1* is an Arf-directed GTPase-activating protein (GAP) shown to modulate actin cytoskeletal dynamics by regulating Arf and Rho family members. We have previously shown that *Arap1^-/-^* mice develop photoreceptor degeneration similar to the human condition retinitis pigmentosa (RP), corroborated by fundus examination, histopathology, and ERG analysis. However, *Arap1* expression was not detected in photoreceptors, but in Müller Glia and retinal pigment epithelium (RPE), suggesting a non-cell-autonomous mechanism for degeneration. The aim of this study was to elucidate the role of retinal *Arap1* in photoreceptor maintenance.

**Methods:** Albino *Arap1^-/-^* mice were generated via breeding pigmented *Arap1^-/-^* mice onto a *Tyr^-/-^* C57BL/6J background. Conditional knockout (cKO) mice were generated for Müller Glia/RPE, Müller Glia, and RPE via targeting *Cralbp, Glast*, and *Vmd2* promoters, respectively, to drive *Cre* recombinase expression to knock out *Arap1*. Mice were analyzed by fundus photography, optical coherence tomography (OCT), histology, and immunohistochemistry. Arap1 binding partners were assayed by affinity purification mass spectrometry.

**Results:** *Vmd2-Cre Arap1^tm1c/tm1c^* and *Cralbp-Cre Arap1^tm1c/tm1c^* mice, but not *Glast-Cre Arap1^tm1c/tm1c^* mice, recapitulated the photoreceptor degeneration phenotype originally observed in germline *Arap1^−/−^* mice. These findings were corroborated by fundus exam, OCT, and histological analysis. Mass spectrometry analysis of ARAP1 co-immunoprecipitation identified putative binding partners of ARAP1, revealing numerous interactants involved in phagocytosis, cytoskeletal composition, intracellular trafficking, and endocytosis. Quantification of rod outer segment (OS) phagocytosis in vivo demonstrated a clear phagocytic defect in *Arap1^−/−^* mice compared to *Arap1^+/+^* littermate controls while cone phagocytosis was preserved.

**Conclusions:** *Arap1* expression, specifically in RPE, is necessary for photoreceptor survival due to its indispensable function in RPE phagocytosis. We propose a model in which Arap1 regulates G-protein function for nonmuscle myosin II targeting during phagocytosis. This novel role of *Arap1* is important for further understanding of both the diversity of its functions and the complex molecular regulation of RPE phagocytosis.

## INTRODUCTION

Retinitis pigmentosa is a rod/cone dystrophy characterized by progressive photoreceptor loss accompanied by striking pigmentary changes on fundus exam.^1^ Generally, cell death follows a progression of early rod loss followed by cone degeneration.^1^ Currently, RP is the leading cause of heritable blindness, with over 60 gene mutations, 35 of which are autosomal recessive, identified for its nonsyndromic form.^2^

Previously we discovered that *Arap1^-/-^* mice, generated by the National Institutes of Health Knockout Mouse Production and Phenotyping (KOMP2) Project, develop a phenotype similar to human RP.^3^ *Arap1* is an Arf-directed GAP with a RhoGAP and multiple Pleckstrin homology (PH) domains. As such, *Arap1* governs a diverse range of functions. *Arap1* has been shown to facilitate EGFR endocytosis and consequent regulation of EGFR signal transduction.^4^ Additionally, *Arap1* has been shown to play a key role in actin modulation as an intersecting node for both Arf- and Rho-directed actin dynamics.^5^ However, *Arap1* function in the retina had not yet been assessed prior to our investigations.

*Arap1^-/-^* mice demonstrated optic nerve pallor, attenuated retinal arteries, retinal pigmentary changes, and focal areas of RPE atrophy, a constellation of findings consistent with those seen in human RP. Optical coherence tomography (OCT) and histopathology revealed outer retinal degradation, most marked in the outer nuclear layer (ONL), with inner retina preservation. These changes were corroborated by reduced scotopic responses and later photopic signal reduction on ERG. As we investigated mechanistic explanations for this phenotype, we found that *Arap1^-/-^* mice experienced normal retinal histogenesis, though soon followed by progressive photoreceptor loss starting with rods. Despite the photoreceptor degeneration observed in the knockout, *Arap1* was not found to be expressed in the photoreceptors, but rather in the adjacent support cells: Müller glia and retinal pigmented epithelium (RPE). In our initial study, due to their pigmented nature, *Arap1* expression was difficult to assess in the RPE.^3^

Müller glia are retinal cells that govern multiple essential functions in retinal homeostasis, including maintenance of retinal structural integrity, glucose metabolism, and neurotransmitter reuptake.^6^ The RPE are a monolayer of cells located beneath the photoreceptor layer known to manage the transport of nutrients, ions, and water, protect against photooxidation, reisomerize all-trans-retinal into 11-cis-retinal, secrete essential factors for retinal maintenance, and phagocytose photoreceptor outer segments.^7^ Dysfunction in both cell types has been implicated the development and pathology of RP. For instance, *MERTK*, encoding a receptor tyrosine kinase involved in RPE phagocytosis, and *RPE65*, encoding an isomerohydrolase essential for visual pigment recycling, have both been linked as RP-causative mutations in the RPE.^8,9^ Reactive gliosis governed by Müller glia has been described extensively in retinitis pigmentosa models, though not many single gene disorders of the retina are directly linked to Muller glial specific genes.^10^ To date, *Arap1* has not been established to play a role in photoreceptor survival in either of these two cell types. Our current investigations seek to elucidate the mechanistic link between *Arap1* loss in Muller glia and/or RPE cells and subsequent development of an RP-like phenotype in mice.

## MATERIALS AND METHODS

### Animals

#### Generation of Arap1 *Tyr*^*c*-2*J*^/*Tyr*^*c*-2*J*^ Mice

Arap1 germline knockout mice were generated by the U.C. Davis Mouse Biology Program as previously described.^3^ B6(Cg)-Tyrc-2J/J mice were acquired from The Jackson Laboratory (stock # 000058) and bred to Arap1^-/-^ germline knockouts. The progeny were then bred with B6(Cg)-Tyrc-2J/J to obtain Arap1^+/-^ *Tyr*^*c*-2*J*^/*Tyr*^*c*-2*J*^ which were used as founders for this strain, colloquially referred to as albino Arap1 germline knockouts.

#### Generation of Arap1 Condition Mouse Strains

Floxed Arap1 conditional mice on the C57BL/6N background were generated by the U.C. Davis Mouse Biology Program. These founders were bred into C57BL/6J and colony founders were confirmed to be free from the *crb1* mutation (*rd8*) by PCR—all mice in this study were confirmed to be free of *rd8*. Three independent strains were then established for the conditional knockout of Arap1 using established Cre lines: Tg(Slc1a3-cre/ERT)1Nat (referred to as Glast-Cre from The Jackson Laboratory Stock #012586); Tg(BEST1-cre)1Jdun (referred to as Vmd2-cre from The Jackson Laboratory Stock # 017557) and Tg(Cralbp-Cre/ERT) (referred to as Cralbp-Cre that was graciously provided by Dr. Edward Levine). For each colony males Arap1^tm1c/+^ Cre+ were bred with females Arap1^tm1c/tm1c^ to obtain progeny for our study (Arap1^tm1c/+^ Cre+ and Arap1^tm1c/tm1c^ Cre+). For both Glast-Cre and Cralbp-Cre, Cre activity was induced with intraperitoneal injection of tamoxifen (75mg/kg of body weight in corn oil) once per day for five consecutive days beginning from P30.

### Histology

For cryoembedding, eyes were enucleated, promptly fixed with 4% PFA for an hour at ambient temperature and then stepwise dehydrated in 10% sucrose, 20% sucrose and 30% sucrose in PBS. The eyes were embedded in Tissue Plus OCT compound (Fisher Scientific) then snap frozen with on a dry ice-ethanol bath. 12 μm thick sections were then obtained using a cryostat (LEICA CM3050; Leica, Wetzlar, Hesse, Germany).

For paraffin embedding, eyes were processed as described in Sun 2015.^11^ Briefly, eyes were enucleated and snap frozen in dry ice cooled propane. The eyes were then step wise fixed with 3% methanol in 97% methanol by storing in −80°C for 7 days, −20°C overnight, and finally in ambient temperature for 2 days. The eyes were then processed into paraffin beginning by dehydration in 100% ethanol (two changes of 100% ethanol; one hour each), replacement in xylene (2 changes; 15 min each) and then 60°C paraffin (three changes; 1 hour each). 5μm sections were then obtained using a Leica RM2125Rt microtome.

### Immunohistochemistry

Cryosections were washed in 1x PBS (3 times; 5 minutes each) and blocked for 1 hour at ambient temperature with blocking solution (4% BSA in 10mM Tris-Cl pH 7.4, 10mM MgCl2, and 0.5% v/v Tween 20). Primary antibodies were then diluted in blocking buffer as specified and applied to the sections overnight at 4°C. Before corresponding Alexa Fluor plus secondary antibodies were added for 1 hour at ambient temperature, the cryosections were washed in PBS (3 times; 5 minutes each). 1μg/ml DAPI was added to the cryosections at ambient temperature for 5 minutes and then the slides were washed in PBS (3 times; 5 minutes each). Cryosection were then coverslipped with FluorSave Reagent (Millipore).

For paraffin sections, sections were deparaffinized using xylenes (3 times; 5 minutes each). The tissue sections were then stepwise hydrated by submersion in 100% ethanol (2 times; 5 minutes each), 95% ethanol (5 minutes), 75% ethanol (5 minutes), and distilled water. Heat-induced epitope retrieval was then performed with 1mM EDTA, pH 8.0. The tissue sections were then blocked for 30min in blocking solution in ambient temperature. Similarly, primary antibodies, DAPI and cover slipping was done as described in the cryosection methods.

### TUNEL

Detection of apoptotic cells was determined by using ApopTag® Fluorescein In Situ Apoptosis Detection Kit (Sigma Aldrich). Paraffin sections were deparaffinized and hydrated as described and then treated with proteinase K (20 μg/mL) at ambient temperature. The tissue sections were then washed with PBS (2 times; 2 minutes each). The tissue sections were equilibrated and TdT enzyme was applied for an hour at 37°C. The reaction was stopped with stop buffer and the sections were washed in PBS (3 times; 1 minute each). The antidigoxigenin conjugate was applied for 30 minutes at ambient temperature followed by a DAPI and cover slipping as described. Finally, the slides were cover slipped with FluorSave Reagent (EMD Millipore). TdT enzyme, equilibration buffer, stop buffer and anti-digoxigenin conjugate were provided with the kit.

### Hematoxylin and Eosin Staining

Sections were deparaffinized using xylenes (3 times; 5 minutes each). The tissue sections were then stepwise hydrated by submersion in 100% ethanol (2 times; 5 minutes each), 95% ethanol (5 minutes), 75% ethanol (5 minutes), and distilled water (5 minutes). The sections were then stained with hematoxylin for 10 minutes, stained with eosin for 5 minutes and dehydrated in reverse of the hydration procedure described beginning from 75% ethanol. The sections were coverslipped with VectaShield (Vector Laboratories).

### B-galactosidase Histochemistry

Cryosections were washed in 1x PBS (3 times; 5 minutes each) and incubated in X-gal containing solution (1 mg/ml X-gal, 5mM potassium ferricyanide, 5mM potassium ferrocyanide and 2 mM MgCl2 in 1x PBS) for 24 hours in a 37°C incubator. The cryosections were then washed in 1x PBS (3 times; 5minutes each) and coverslipped with VectaShield.

### Ocular Imaging: Fundus Photographs and Optical Coherence Tomography

Mice were sedated with IP injections with a combination of ketamine (50mg/kg) and dexmedetomidine (0.25-0.5 mg/kg). The eyes were dilated with 2.5% phenylephrine hydrochloride (Akorn, Lake Forest, IL, USA) and tropicamide (Bausch & Lomb, Tampa, FL, USA); the eyes were lubricated with GenTeal (Alcon Laboratories, Fort Worth, TX, USA). Fundus images were obtained with StreamPix 5 using a Micron III (Phoenix Research Laboratories, Pleasanton, CA, USA).

Optical coherence tomography was acquired using Envisu R2200 SDOIS Imaging System (Bioptigen, Morrisville, NC, USA) and analyzed using InVivoVue 2.4.35.

### Scanning Electron Microscopy

Eyes were fixed in 2.5% glutaraldehyde and 2% paraformaldehyde in 0.1M sodium cacodylate buffer (all from Electron Microscopy Sciences) overnight with gentle rocking at 4°C, washed with 0.1 M cacodylate buffer, and post-fixed in 1% osmium tetroxide for 2 h at ambient temperature. The eyes were then dehydrated in a graded ethanol series, further dehydrated in propylene oxide and embedded in Epon epoxy resin. Semi-thin (1 μm) and ultra-thin sections were cut with a Leica EM UC6 ultramicrotome and the latter were collected on pioloform-coated (Ted Pella) one-hole slot grids. Sections were contrasted with Reynolds lead citrate and 8% uranyl acetate in 50% ethanol and imaged on a Philips CM120 electron microscope equipped with an AMT BioSprint side-mounted digital camera and AMT Capture Engine software.

### Cell Culture

Discarded de-identified human fetal eyes were collected for use in cell culture with permission of the UC Davis IRB. Eyes were carefully dissected by removing the anterior segment, carefully removing the retina, and peeling the retinal pigmented epithelium as a single sheet (monolayer) with the use of a dissecting microscope. Fetal retinal pigment epithelia were cultivated at 37°C under a humidified 5% CO2 atmosphere in 3D-Retinal Differentiation Media (3:1 Dulbecco’s modified Eagle medium (DMEM)-F12, DMEM (High Glucose) supplemented with 5% (v/v) Fetal Bovine Serum (Atlanta Biologicals), 2 mM GlutaMAX (Thermo Fisher), 200 μM Taurine, 1x Penicillin/Streptomycin (Thermo Fisher), 1:1000 chemically defined lipid supplement (Thermo Fisher), 2% B27 supplement (Thermo Fisher) as described previously.^12^

### Protein Extraction

Differentiated cells were washed with ice cold 1x PBS and incubated on ice for 10 minutes in ice cold lysis buffer (25 mM Tris-HCl pH 7.4, 150 mM NaCl, 1 mM EDTA, 1% NP-40, 5% glycerol, and protease inhibitors). Cells were then mechanically detached with a cell scraper and homogenized via Dounce homogenizer. For mouse liver samples, 250 μg of liver was homogenized in 750 μL of lysis buffer via Dounce homogenizer. Homogenate was centrifuged at 4°C at 14,000 x g for 20 minutes and supernatant was collected.

### Immunoprecipitation

Lysates were precleared on Protein G-magnetic Beads (Thermo Fisher) for 1 hour at 4°C. Protein G-beads were incubated with antibodies for 1 hour and subsequently cross-linked with 5 mM BS3 for 30 minutes at ambient temperature. Immunoprecipitation was performed per Dynabeads™ Protein G Immunoprecipitation Kit (Thermo Fisher) with goat anti-Arap1 (1:50, Abcam) and normal goat IgG (1:50, R&D Systems).

### Western Blot

Lysates and immunoprecipitates were loaded onto NuPAGE™ 4 to 12%, Bis-Tris, 1.5 mm gels (Thermo Fisher) and resolved. Gels were wet-blotted onto polyvinylidene fluoride (PVDF) membranes and subsequently blocked with 5% BSA in TBST for 1 hour at ambient temperature. Blots were incubated with goat anti-Arap1 IgG (1:2000, Abcam) and rabbit anti-Beta-actin IgG (1:4000, Cell Signaling) overnight at 4°C. After incubation with HRP-conjugated donkey anti-goat IgG (1:4000, Abcam) and HRP-conjugated goat anti-rabbit IgG (1:4000, Cell Signaling) for 1 hour at ambient temperature, bands were detected with enhanced chemiluminescence Western blotting detection reagents (Amersham).

### Sample Preparation

Protein from each immunoprecipitated were subjected to tryptic digestion via suspension-trap (S-Trap) devices (ProtiFi). S-Trap is a powerful Filter-Aided sample preparation (FASP) method that consists in trapping acid aggregated proteins in a quartz filter prior enzymatic proteolysis, and allows for reduction / alkylation / tryptic proteolysis all in one vessel. Specifically, proteins were resuspended in 50 μL Solubilization Buffer. Solubilization Buffer consists of 5% SDS, 50 mM triethyl ammonium bicarbonate, complete protease inhibitor cocktail (Roche), pH 7.5 Disulfide bonds were reduced with dithiothreitol and alkylated with iodoacetamide in 50mM TEAB buffer. The enzymatic digestion was a first addition of trypsin 1:100 enzyme: protein (wt/wt) for 4 hours at 37°C, followed by a boost addition of trypsin using same wt/wt ratios for overnight digestion at 37°C. Peptides were eluted from S-Trap by sequential elution buffers of 100mM TEAB, 0.5% formic acid, and 50% acetonitrile 0.1% formic acid. The eluted tryptic peptides were dried in a vacuum centrifuge and re-constituted in 0.1% trifluoroacetic acid. These were subjected to LC-MS analysis.

### LCMS

Peptides were resolved on a Thermo Scientific Dionex UltiMate 3000 RSLC system using a PepMap 75μmx25cm C18 column with 2 μm particle size (100 Å pores), heated to 40°C. A 0.6 μg of total peptide amount was injected for each sample, and separation was performed in a total run time of 90 min with a flow rate of 200 μL/min with mobile phases A: water/0.1% formic acid and B: 80%ACN/0.1% formic acid. Gradient elution was performed from 10% to 8% B over 3 min, from 8% to 46% B over 66 min, and from 46 to 99% B over 3 min, and after holding at 99% B for 2 min, down to 2% B in 0.5 min followed by equilibration for 15min. Peptides were directly eluted into an Orbitrap Exploris 480 instrument (Thermo Fisher Scientific, Bremen, Germany). Spray voltage were set to 1.8 kV, funnel RF level at 45, and heated capillary temperature at 275 °C. The full MS resolution was set to 60,000 at m/z 200 and full MS AGC target was 300% with an IT set to Auto. Mass range was set to 350–1500. AGC target value for fragment spectra was set at 200% with a resolution of 15,000 and injection time was set to Standard and Top40. Intensity threshold was kept at 5E3. Isolation width was set at 1.6 m/z, normalized collision energy was set at 30%.

### Raw Data Processing

The LCMS .raw files were processed with Proteome Discoverer 2.4 (Thermo Fisher) using the integrated SEQUEST engine. All data was searched against a target/decoy version of the human Uniprot Reference Proteome without isoforms (21,074 entries). Peptide tolerance was set to 10 ppm, and fragment mass tolerance was set to 0.6 Da. Trypsin was specified as enzyme, cleaving after all lysine and arginine residues and allowing up to two missed cleavages. Carbamidomethylation of cysteine was specified as fixed modification and protein N-terminal acetylation, oxidation of methionine, deamidation of asparagine and glutamine and pyro-glutamate formation from glutamine were considered variable modifications with a total of 2 variable modifications per peptide. The false discovery rate limit of 1-5% on the peptide level.

### In Situ Phagosome Quantification

Retinal cryosections were washed as described above and subsequently blocked at ambient temperature for 1 hour. Sections were then incubated with mouse anti-rhodopsin IgG (1:1000, EMD Millipore) and rabbit anti-L/M opsin IgG (1:1000, EMD Millipore) overnight at 4°C. Sections were washed with 1x PBS (3 times; 5minutes each) and incubated with donkey anti-mouse Alexa Fluor Plus 488 IgG (1:500, Thermo Fisher) and donkey anti-rabbit Alexa Fluor Plus 647 IgG (1:500, Thermo Fisher) at 37°C for 1 hour. Slides were then stained for DAPI and washed as described above. Sections were coverslipped with FluorSave Reagent and imaged with an Olympus FV3000 Confocal Laser Scanning Microscope (Olympus Corporation, Shinjuku City, Tokyo, Japan). Rhodopsin- and cone opsin-positive phagosomes were quantified as previously described.^13,14^

## RESULTS

### Loss of *Arap1*, expressed in Müller glia and RPE, is associated with photoreceptor death

Previously, we reported photoreceptor degeneration in *Arap1^-/-^* mice on a pigmented C57BL/6J background. Using the LacZ cassette knocked into the *Arap1* locus under control of its promoter, we were able to use an X-gal reaction to detect *Arap1* expression in the retina. However, pigment in the RPE layer made X-gal signal detection and subsequent verification of *Arap1* expression difficult. To better assess RPE expression of *Arap1*, we bred *Arap1^-/-^* mice onto a *Tyr^-/-^* C57BL/6J albino background. To confirm that the pattern of retinal degeneration seen originally in the pigmented *Arap1^-/-^* mice was still present, we sectioned eyes and stained with hematoxylin and eosin to visualize retinal morphology. Albino *Arap1^-/-^* mice exhibited thinning of the ONL and with preservation of inner retinal morphology (Fig. 1A), identical to pigmented *Arap1^-/-^* mice described previously.^3^ These changes were absent in albino *Arap1^+/+^* littermates (Fig. 1B).

**Figure 1.**
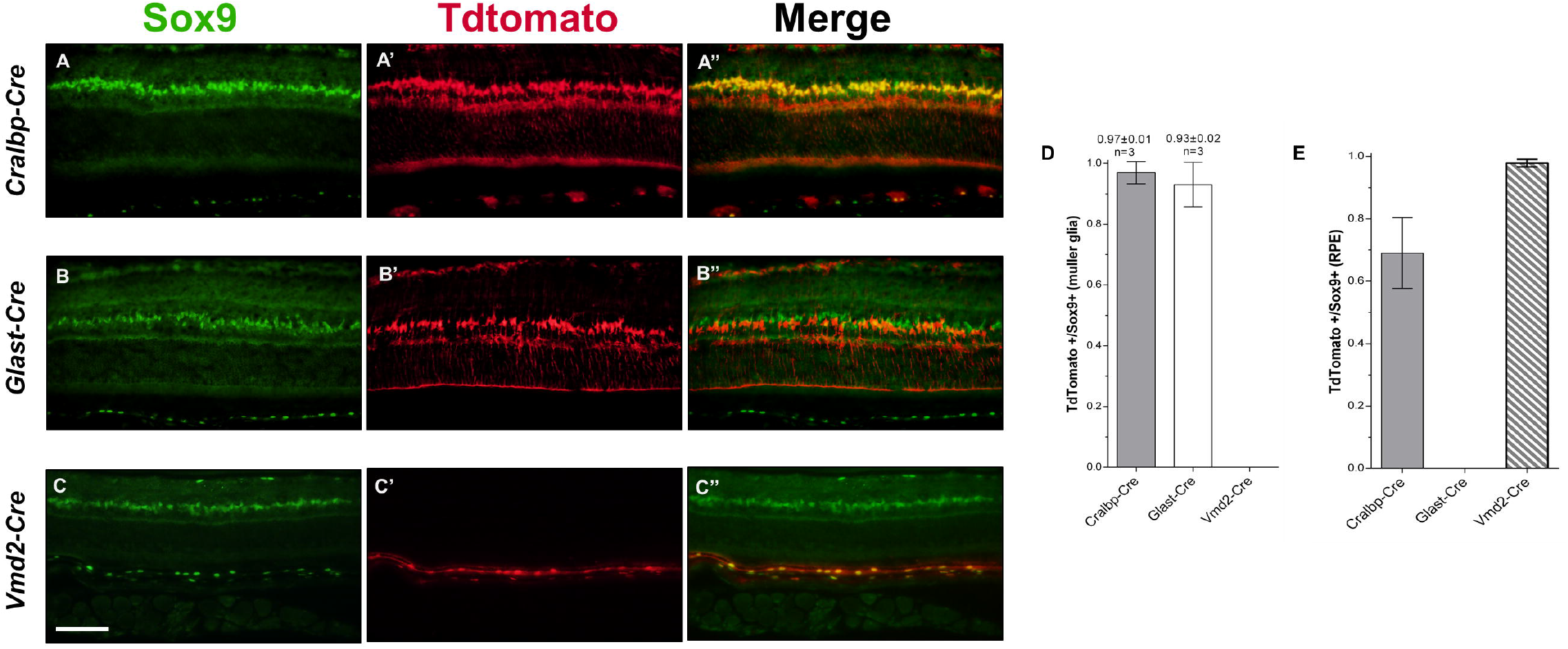
Characterization of Albino Arap1^-/-^ Retina. Hematoxylin-eosin staining of representative retinal sections of (**A**) *Arap1^-/-^* pigmented (top) and albino (bottom) and (**B**) *Arap1^+/+^* pigmented (top) and albino (bottom) are shown. Retinal thinning and photoreceptor degeneration is prominent in the pigmented and albino *Arap1^-/-^* retinas but not seen in *Arap1^+/+^* retinas. Representative retinal sections of albino *Arap1^-/-^* (**C**) and *Arap1^+/+^*(**D**) were used for terminal deoxynucleotidyl transferase end labelling assay (TUNEL, green) and immunohistochemistry for cleaved PARP-1 (cPARP, red) fluorescent staining to assess programmed cell death. DAPI counterstaining (blue) was used to visualize the nuclei of the retinal layers. Both cPARP and TUNEL signals were detected in the ONL of *Arap1^-/-^* retinas but not in control *Arap1^+/+^* retinas. Retinal sections of *Arap1^+/-^* mice were stained with X-gal to assess LacZ histochemical reaction (**E**). Blue X-gal signal was detected in the INL, ONL, and RPE (arrows) with rare signal in the GCL. cPARP, TUNEL, and X-gal analysis were performed on tissue harvested from mice aged postnatal day 24, while hematoxylin and eosin staining from mice aged postnatal 6 weeks. The ganglion cell layer (GCL), inner plexiform layer (IPL), inner nuclear layer (INL), outer plexiform layer (OPL), outer nuclear layer (ONL), inner segments (IS), outer segments (OS), and retinal pigment epithelium (RPE) are labeled (**B**). All images were taken at 40x magnification. Scale bar represents 100 μm (**B,C**). N ≥ 3 for each group.

To assess if photoreceptor loss was due to programmed cell death, we analyzed apoptotic signal with both terminal deoxynucleotidyl transferase end labelling (TUNEL) and cleaved PARP-1 (cPARP) fluorescent staining. TUNEL staining visualizes DNA fragmentation secondary to apoptosis, while cPARP staining measures caspase-3 activity. Consistent with the pattern of degeneration observed on H&E staining, apoptotic signals were observed in the ONL in both TUNEL and cPARP fluorescence (Fig. 1C). In contrast, apoptotic signals were absent in the ONL in *Arap1^+/+^* control retinas (Fig 1D).

To assess *Arap1* expression in the RPE, we used our albino *Arap1^+/-^* mice and the signal from X-gal histochemical reaction as a surrogate for *Arap1* expression. In postnatal day 24 retina, blue X-gal signal was clearly seen in the RPE and the inner nuclear layer of albino *Arap1^+/-^* mice (Fig. 1E). We confirmed previously that the expression pattern observed in these layers was due to Müller glia.^3^ However, signal was also observed in the RPE layer that was originally difficult to visualize in pigmented *Arap1^+/-^* mice, which is clearly visible on an albino background (Fig. 1E).

### Conditional knockout of *Arap1* in RPE but not Müller glia recapitulates the *Arap1^-/-^* photoreceptor degeneration phenotype

To confirm that Arap1 expression specifically in the retina is essential for photoreceptor viability, particularly in Müller glia and RPE as X-gal reactivity suggested, we generated a tamoxifen-induced Müller glia/RPE-specific conditional *Arap1* knockout mouse. By using the *Cralbp* promoter to drive *Cre* recombinase expression, knockout was ensured to be tissue-specific to Müller glia and RPE.^15^ To assess both efficacy and specificity of targeting, Cre function in *Cralbp-Cre* mice was quantified by breeding these mice onto an Ai9 background to generate a TdTomato signal (red) in cells with Cre activity. Anti-Sox9 (green), a transcription factor present in Müller glia and the RPE, was used to mark these nuclei for quantification of Cre function.^16,17^ TdTomato fluorescence mirrored Sox9 immunosignal, confirming Cre function specific to the RPE and Müller glia (Fig 2A,2A’,2A’’). *Cre* function was present in nearly 100% of Müller glia and ~70% of the RPE (Fig. 2D,2E).

**Figure 2.**
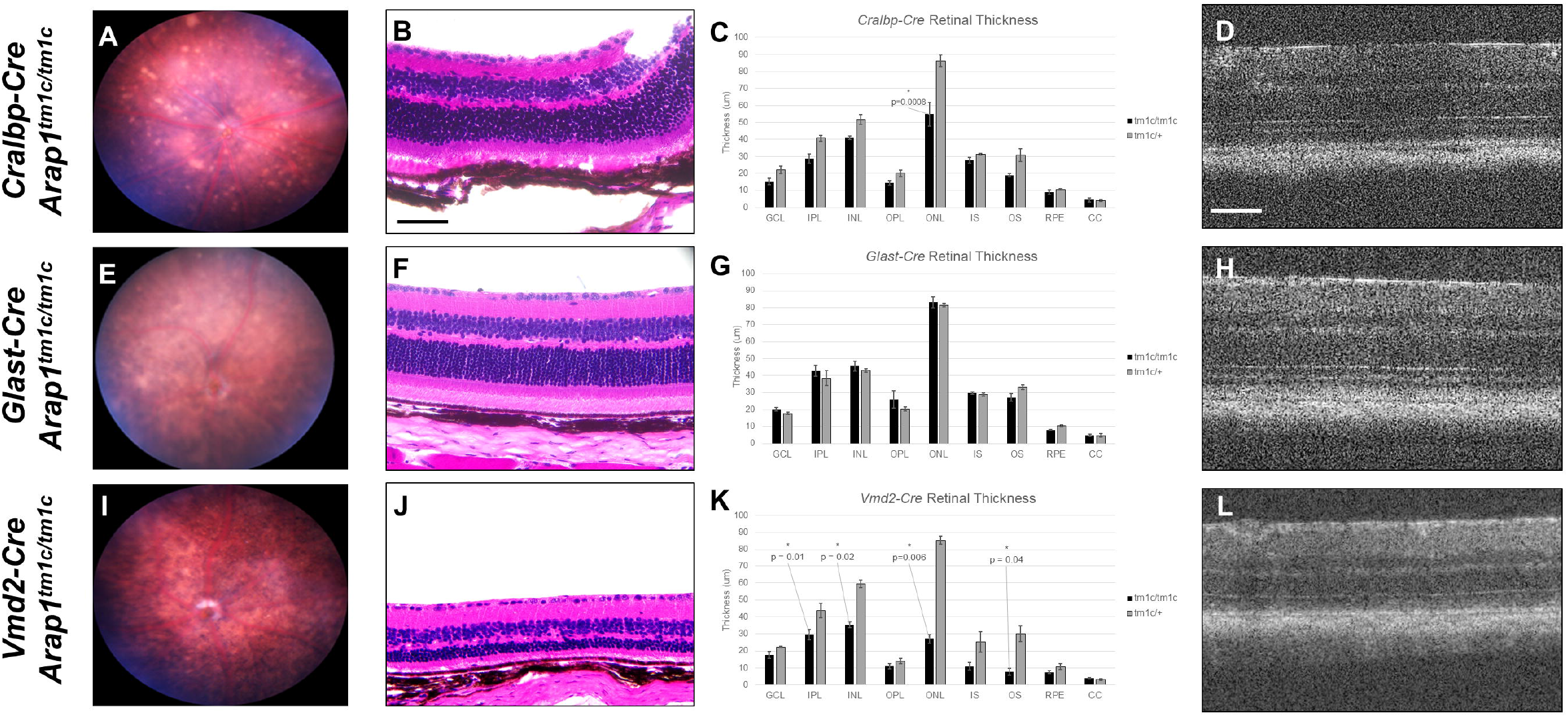
Quantification of Cre Function in Conditional Knockout Mice. Immunohistochemistry was performed using Anti-Sox9 (green) and Anti-Tdtomato (red) to quantify *Cre* function in *Cralbp, Glast*, and *Vmd2-Cre* mice postnatal day 84. Sox9 immunosignal was detected in all of the Muller glia and RPE cells. in all Cre lines (**A,B,C**). Tdtomato signal was visualized in *Cralbp-Cre, Glast-Cre*, and *Vmd2-Cre* mouse lines (**A’,B’,C’**). Channels were merged to create a composite image. Tdtomato signal was present in the Muller Glia in the *Cralbp-Cre* and *Glast-Cre* strains, but not the *Vmd2-Cre* strain. Conversely, Tdtomato signal was present in RPE in the *Cralbp-Cre* and *Vmd2-Cre* strains, but not the *Glast-Cre* strain. Merged Tdtomato/Sox9 images are shown (**A’’,B’’,C’’**) with graphical quantification of the proportion of Sox9-positive cells that were also TdTomato-positive in Muller glia (**D**) and RPE (**E**) for each *Cre* line. Images were taken at 40x magnification. Scale bar represent 100 μm (**C**). N = 3 for each group, error bars represent SE.

*Cralbp-Cre Arap1^tm1c/tm1c^* mice shared phenotypic features that were originally observed in *Arap1^-/-^* mice, though generally milder. Two months post-tamoxifen induction, fundus examination revealed retinal pigmentary changes, focal areas of RPE atrophy, and areas of spotty hyperreflective material (Fig. 3A). Histopathology revealed degenerative loss of the outer nuclear layer as well as degenerative changes of the outer retina (Fig. 3B,3C). These findings were corroborated by OCT imaging revealing thinning of the outer retina with preservation of the inner retina (Fig 3D).

**Figure 3.**
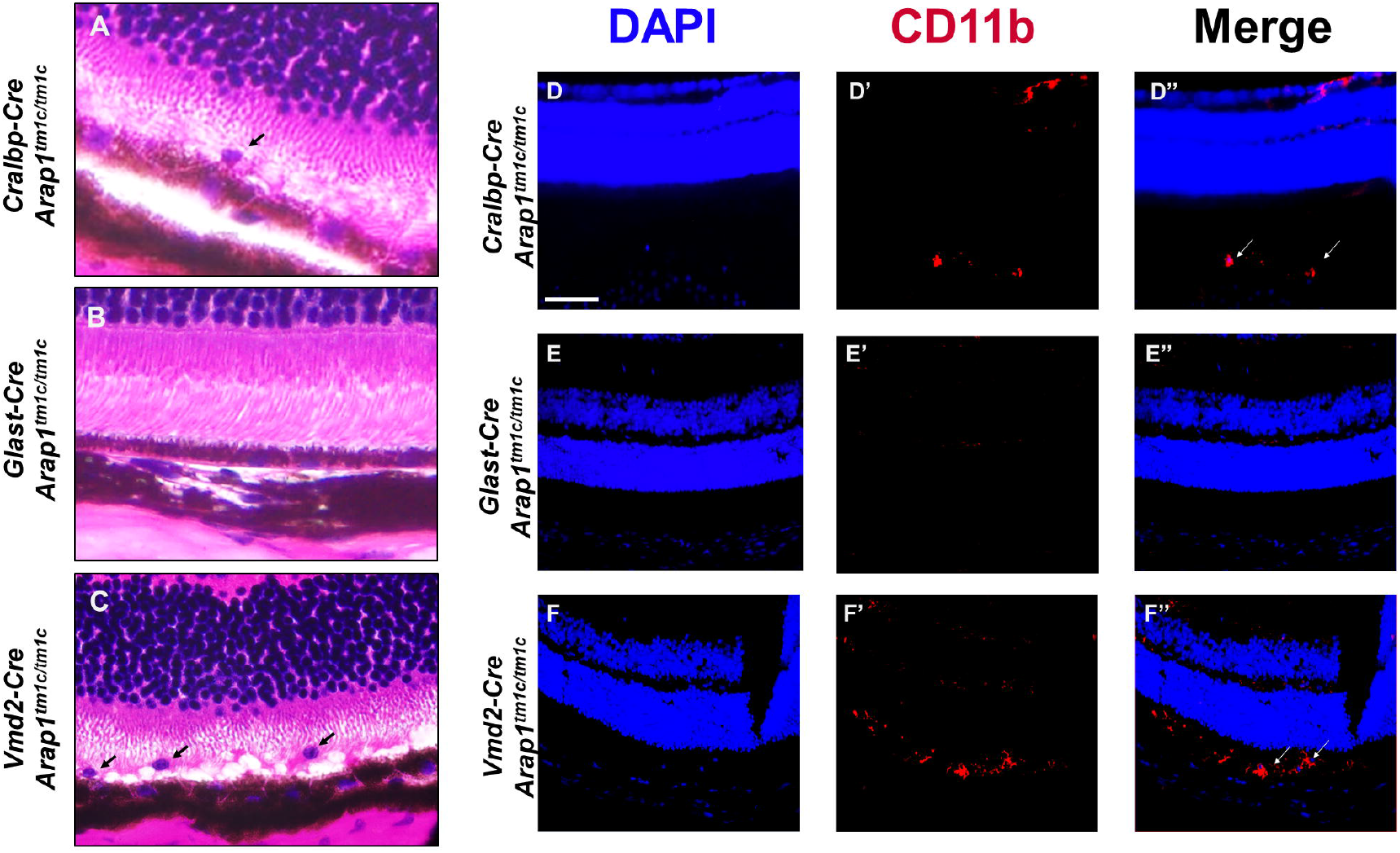
Characterization of Cralbp Arap1^tm1c/tm1c^, Glast Arap1^tm1c/tm1c^, and Vmd2-Cre Arap1^tm1c/tm1c^ Conditional Knockout Mouse Lines. *Cre* cKO mice were analyzed with fundus photography, histology, and OCT analysis at 3 months of age (Glast-Cre, Cralbp-Cre) and 1 month of age (Vmd2-Cre). Fundus photography demonstrated RPE atrophy, pigmentary changes, optic nerve pallor, and vascular attenuation in the *Crablp-Cre Arap1^tm1c/tm1c^* and *Vmd2-Cre Arap1^tm1c/tm1c^* strains (**A,I**). Conversely, the *Glast-Cre Arap1^tm1c/tm1c^* strain demonstrated no significant differences from wild type littermates (**E**). Representative retinal sections from the *Cre* strains were stained with hematoxylin and eosin. Quantification of retinal layers on histology is shown for each cKO line compared to conditional heterozygote controls (**C,G,K**). *Cralbp-Cre Arap1^tm1c/tm1c^* retinas demonstrated significant degeneration of the ONL with relative preservation of all other retinal layers (**B,C**). *Glast-Cre Arap1^tm1c/tm1c^* retinas were indistinguishable from tm1c/+ littermate retinas (**F,G**). *Vmd2-Cre Arap1^tm1c/tm1c^* retinas demonstrated more severe degeneration of the ONL (**J**) and quantification of retinal layers revealed significant degeneration of the IPL, ONL, and OS layers compared to heterozygous littermates (**K**). These changes were consistent with in vivo OCT imaging of retinal layers (**D,H,L**). Scale bar (**B,D**) represents 100 μm. Images were taken at 40x magnification (**B,F,J**). N=3, error bars represent standard error, p values are shown in graph.

To ascertain whether *Arap1*’s role in Müller glia or RPE was determinant for the *Arap1^-/-^* phenotype, conditional knockouts using *Glast* and *Vmd2* promoters respectively, were generated using the same method as the *Cralbp-Cre* mice. The *Vmd2* promoter is RPE-specific, while the *Glast* promoter is Müller glia-specific.^18,19^ Endogenous fluorescence of TdTomato and immunohistochemistry using anti-Sox9 was used again to quantify the *Cre* function in Muller nd cell patterns in both the *Glast-Cre* and *Vmd2-Cre* lines (Fig. 2B,2B’,2C,2C’). *Glast-Cre* TdTomato signal mirrored Sox9 signal in Müller glia, though lacked any observable signal in RPE (Fig. 2B’’). *Vmd2-Cre* TdTomato signal, contrastingly, was only detected in the RPE layer (Fig. 2C’’). Both knockouts expressed *Cre* in nearly 100% of their respective cells (Fig. 2D,2E), confirming generation of Müller glia and RPE cell type-specific conditional knockouts, respectively.

Despite the significant degree of *Arap1* expression in Müller glia, *Glast-Cre Arap1^tm1c/tm1c^* mice were phenotypically indistinguishable from wild type mice. Fundus exam, histochemistry, and OCT analysis revealed no significant changes from heterozygous littermates (Fig 3E-H). In contrast, *Vmd2-Cre Arap1^tm1c/tm1c^* mice recapitulated the phenotype observed in *Cralbp-Cre Arap1^tm1c/tm1c^* and *Arap1^-/-^* mice (Fig 3I-L). Fundus exam again revealed optic nerve pallor, attenuated retinal arteries, retinal pigmentary changes, generalized RPE atrophy, and outer retinal thinning on histology and OCT. Histopathology was similar to *Arap1^-/-^* mice and more severe than *Cralbp-Cre Arap1^tm1c/tm1c^* mice with significant degeneration of the IPL, INL, ONL, and OS compared to heterozygous littermates (Fig 3J).

Hematoxylin and eosin staining revealed invasion of cells in the outer retina of *Vmd2-Cre Arap1^tm1c/tm1c^* and *Cralbp-Cre Arap1^tm1c/tm1c^* retinas that were suspicious for macrophages, absent in *Glast-Cre Arap1^tm1c/tm1c^* retinas (Fig. 4A-C). To confirm this finding, immunohistochemistry was performed using anti-CD11b to visualize retinal macrophages. Analysis revealed CD11b signal in the outer retina of *Cralbp-Cre Arap1^tm1c/tm1c^* and *Vmd2-Cre Arap1^tm1c/tm1c^* mice, indicative of macrophage invasion (Fig. 4D’’,4F’’). *Glast-Cre Arap1^tm1c/tm1c^* mice did not demonstrate any measurable CD11b signal in their retinas (Fig. 4E’’).

**Figure 4.**
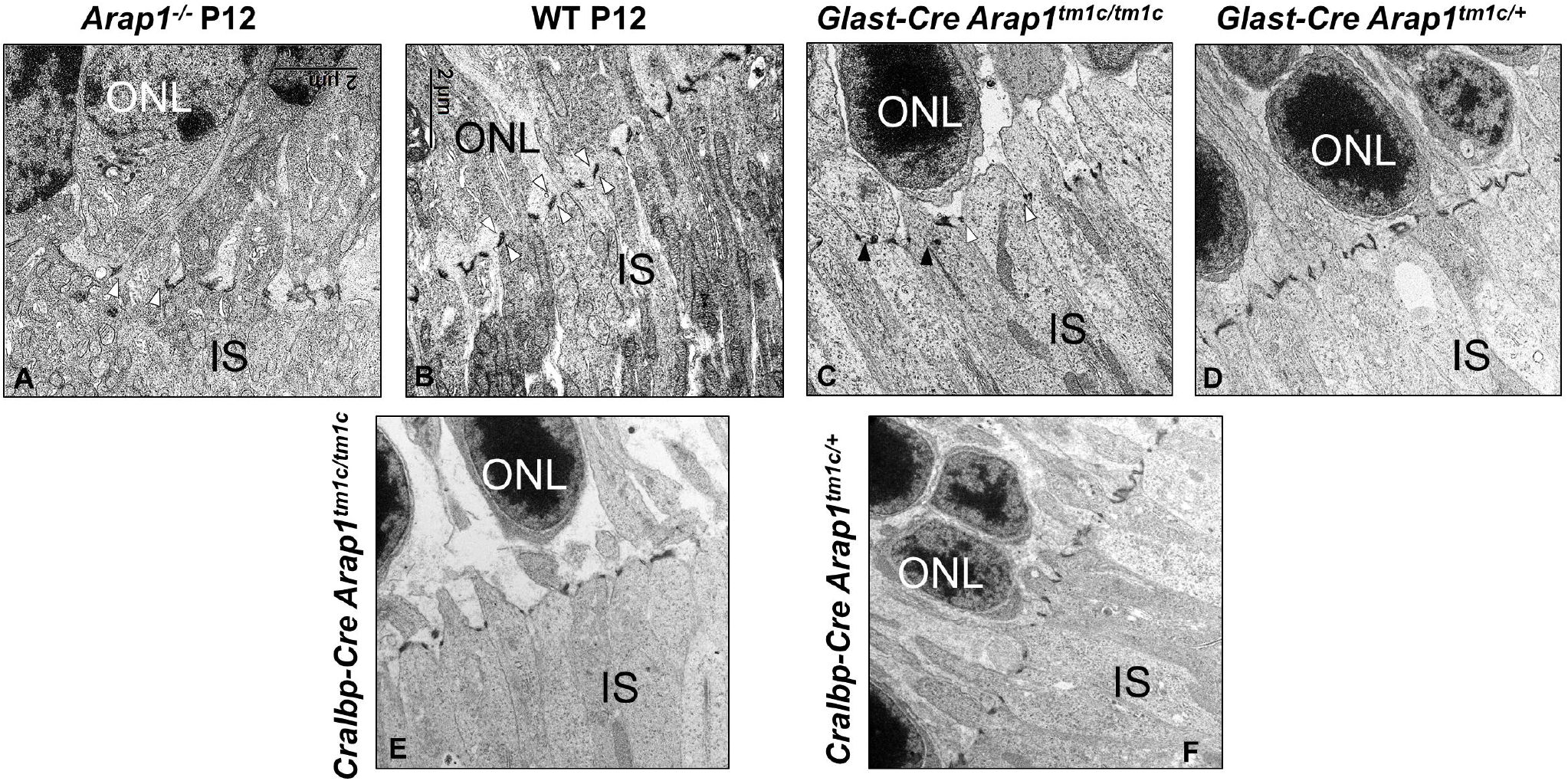
Outer Retinal Macrophage Invasion in Cre cKO mice. Hematoxylin and eosin staining revealed cells suspicious for macrophages in the outer retina of *Vmd2-Cre Arap1^tm1c/tm1c^* and *Cralbp-Cre Arap1^tm1c/tm1c^* mice (**A,C**; arrows), but absent in *Glast-Cre Arap1^tm1c/tm1c^* retinas (**B**). To confirm this finding, immunohistochemistry was performed with anti-CD11b (red) (**D’,E’,F’**) with DAPI (blue) (**D,E,F**) counterstaining to visualize the nuclei of the retinal layers in animals aged postnatal day 84. Channels were merged to create a composite image (**D’’,E’’,F’’**). CD11b signal was detected in the outer retina of *Vmd2-Cre Arap1^tm1c/tm1c^* and *Cralbp-Cre Arap1^tm1c/tm1c^* mice (**D’’,F’’**; arrows), indicative of macrophage invasion. *Glast-Cre Arap1^tm1c/tm1c^* retinas lacked any significant signal (**E’’**). Images were taken at 40X magnification; scale bar (**D**) represents 100 μm.

Despite the lack of any discernable phenotype by retinal imaging and histology, *Glast-Cre Arap1^tm1c/tm1c^* retinas have structural abnormalities of Muller glia, as detected by transmission electron microscopy. Compared to wild type retinas (Fig. 5B), the external limiting membrane (ELM) of *Arap1^-/-^, Glast-Cre Arap1^tm1c/tm1c^*, and *Cralbp-Cre Arap1^tm1c/tm1c^* retinas were discontinuous and disorganized (Fig. 5A,5C,5E). Gaps between adherens junction (AJ) complexes and loss of linear arrangement were common in both germline and conditional mutant retinas. Littermate *Glast-Cre Arap1^tm1c/+^* and *Cralbp-Cre Arap1^tm1c/+^* retinas demonstrated structurally normal ELM (Fig. 5D,5F).

**Figure 5.**
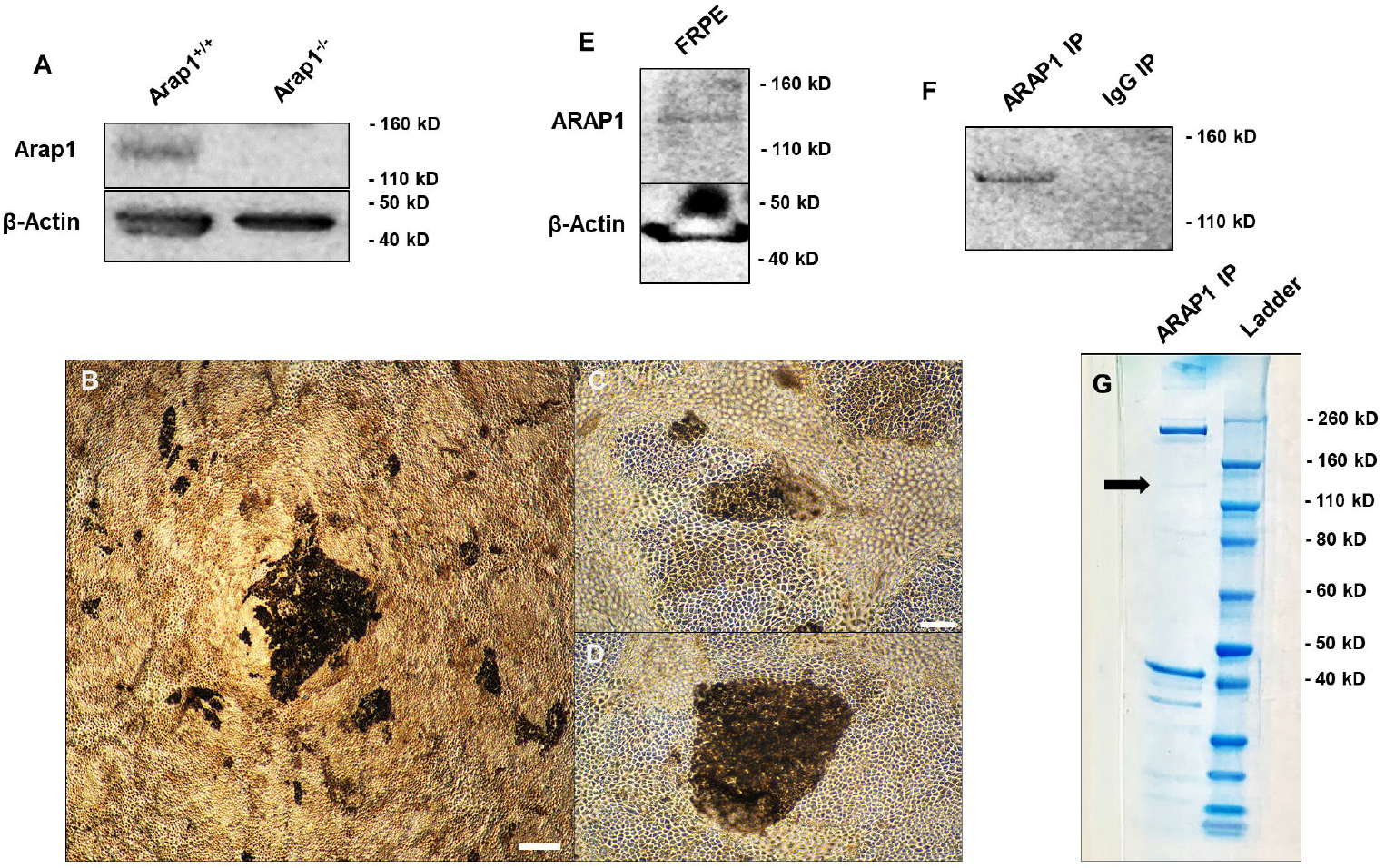
External Limiting Membrane Degeneration in Germline and Conditional Arap1 Knockout Retinas. Transmission electron microscopy (TEM) of representative retinal sections were assessed in wild type, *Arap1^-/-^, Glast-Cre Arap1^tm1c/tm1c^*, and *Cralbp-Cre Arap1^tm1c/tm1c^* mice. Wild type and *Arap1^-/-^* retinas were assessed at postnatal day 12. -*Cre Arap1^tm1c/tm1c^* and -*Cre Arap1^tm1c/+^* retinas were assessed at 8 months of age. *Arap1^-/-^* retinas demonstrate fewer adherens junction (AJ) complexes with increased space between each junction (delineated by the arrowheads) as well as loss of their linear morphology and arrangement (**A**). Wild type retinas demonstrated normal architecture of AJ complexes (arrowheads) with typical linear arrangement (**B**). *Glast-Cre Arap1^tm1c/tm1c^* retinas also demonstrated frequent gaps between AJ complexes (delineated by arrowheads) alternating with regions of normal AJ complexes (black arrowheads) and loss of linear arrangement (**C**). Littermate *Glast-Cre Arap1^tm1c/+^* retinas did not demonstrate abnormalities in the ELM (**D**). *Cralbp-Cre Arap1^tm1c/tm1c^* retinas also demonstrated abnormal ELM junction structure with significant gaps between AJ, though linear arrangement was relatively preserved (**E**). These abnormalities are not observed in *Cralbp-Cre Arap1^tm1c/+^* littermate retinas (**F**). Images were taken at E4300x magnification.

Though not documented in *Crablp-Cre* or *Glast-Cre* mice, *Cre* toxicity in the noninduced *Vmd2-Cre* line has been debated.^20,21^ To account for potential *Cre* cytotoxicity in all cKO lines, we analyzed *Cralbp-Cre Arap1^tm1c/+^, Glast-Cre Arap1^tm1c/+^*, and *Vmd2-Cre Arap1^tm1c/+^* littermate retinas with fundus imaging, histology, and OCT measurement. None of the *Cre* heterozygotes demonstrated significant differences in fundus appearance, histological analysis, and OCT analysis from wild-type mice (Supplemental figure 1). Furthermore, analysis of *Vmd2-Cre Arap1^tm1c/tm1c^* mice was carried out at 1 month of age as no *Cre*-related degeneration has been documented by this time point.^20^

### Human Retinal ARAP1 Interactome

After determining Arap1 is essential in RPE cells for photoreceptor survival, we sought to understand the specific function of this protein in these cells. To elucidate potential cellular processes in which Arap1 is involved, co-immunoprecipitation of ARAP1 and its binding partners was performed on cultured human fetal RPE (FRPE) cells. The antibody, previously validated for western blot detection and immunoprecipitation of ARAP1, was able to detect Arap1 in WT mouse liver lysate, but not *Arap1^-/-^* liver lysate.^22^ A band at approximately 136 kD was observed, consistent with Arap1 isoform 3 (Fig. 6A). Additional bands were observed in the WT lysates potentially belonging to other Arap1 isoforms or non-specific binding (Supp. Fig. 2A). To ensure that this antibody bound human ARAP1, we obtained fetal donor eyes and meticulously dissected RPE tissue for culture. To ensure that protein expression patterns were similar to *in vivo* RPE, cultures were maintained until they adopted physical characteristics of maturity as defined previously by pigmentation and “cobblestone” hexagonal morphology (Fig. 6B-D).^23,24^ Western blot analysis was performed on FRPE lysate, which detected a band at the same molecular weight (Fig. 6E).

**Figure 6.**
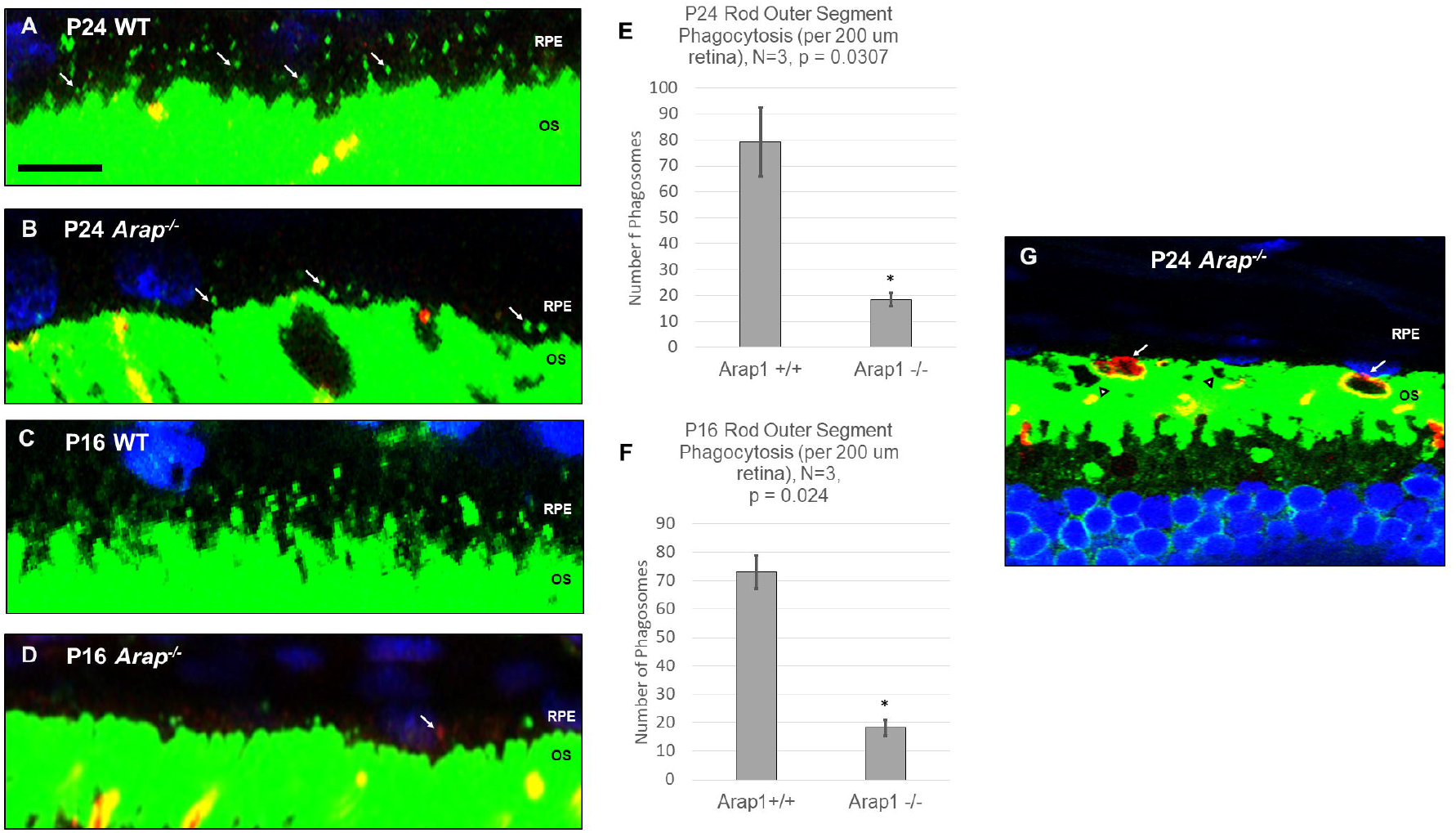
ARAP1 Co-Immunoprecipitation. Western blot analysis validated that the anti-Arap1 antibody detected a band at ~136 kD in *Arap1^+/+^*, but not *Arap1^-/-^* mouse liver lysate (**A**). β-actin was included as an endogenous control. FRPE were harvested from donor tissue and grown in culture. Cells were lysed with NP-40 lysis buffer when mature, as defined by pigmentation and hexagonal “cobblestone” morphology (**B,C,D**; pictured passage 1, culture day 98). Immunoprecipitation of FRPE lysate with anti-ARAP1 was performed with a parallel goat IgG control immunoprecipitation. Immunoprecipitates were analyzed with western blot analysis along with FRPE lysate. A band was detected at ~136 kD in both FRPE lysate (**E**) and anti-ARAP1 immunoprecipitate, but not the control goat IGG immunoprecipitate (**F**). Coomassie analysis of the anti-Arap1 immunoprecipitation is shown in (**G**) with a faint blue band at ~136 kD (arrow). Uncropped blots are shown in Supplemental Figure 2. Scale bar represents 300 um (B) and 100 um (**C**). Images were taken at 4X (**B**) and 10X magnification (**C,D**).

To ensure that putative binding partners of ARAP1 detected by mass spectrometry were not precipitated due to non-specific binding to the magnetic Protein G beads, FRPE lysates were pre-cleared with beads and bead-antibody complexes were cross-linked. To control for non-specific binding due to the antibody itself, non-specific goat IgG immunoprecipitations were carried out in parallel to goat anti-ARAP1 immunoprecipitations. On western blot analysis using FRPE cell lysate, ARAP1 was detected at 136 kD in the anti-ARAP1 immunoprecipitate, but not control anti-IgG immunoprecipitate (Fig. 6F). We analyzed the anti-ARAP1 immunoprecipitate post-gel electrophoresis with Coomassie staining to ensure that there was sufficient protein for mass spectrometry analysis. Coomassie analysis yielded a blue band at approximately 136 kD (Fig. 6G).

Upon initial liquid chromatography-tandem mass spectrometry (LC-MS/MS) analysis, we identified 816 proteins with a threshold of 95%. Further stratification was performed with the following criteria: percentage sequence coverage > 4%, number of unique peptides > 3, relative abundance ratio (anti-ARAP1 IP/control IgG IP) > 5, and significant protein detection in minimum 3 of 5 biological replicates. Post-stratification, ~150 putative binding partners remained. LC-MS/MS detected numerous proteins involved in actin cytoskeletal management that have been detected in the past, including members of actin-related protein 2/3 (ARP-2/3) complex family and F-actin capping protein complex.^22^ Novel interactants in the composition of the actin cytoskeleton were identified as well. These included components of microfilaments (ezrin, calponin-3, moesin) and intermediate filaments (desmocollin-1) essential for the formation of the RPE cytoskeleton as well as F-actin regulators (ankycorbin).^25^ Many members of the myosin family were identified (MYO6, MYO7A, MYH9, MYH10).^25^ These results align with ARAP1’s previously established roles in cytoskeletal coordination.^22,26^

Many established components of the RPE phagocytic machinery were identified, such as unconventional myosin 7-a (MYO7A). Mutations in MYO7A have been linked to Usher syndrome, a condition denoted by hearing loss and RP, and defects in RPE phagocytosis of rod OS.^27^ Subunits of V-type proton ATPase (V-ATPase), a lysosome-associated protein, were identified. V-ATPase has been shown to be an essential component of RPE phagocytic machinery and subsequent photoreceptor maintenance.^28,29^ Components and known interactors of non-muscle myosin type II (NMII) were found, such as MYL6B, MYLK, MYH9, MYH10, and CTNNB1. NMII is an essential participant of RPE phagocytic machinery through its interactions with Mer tyrosine kinase (MerTK).^30^ Other identified proteins implicated in phagocytosis, but with no established role in RPE, include members of the coronin family (coronin-1, coronin-2).^31^

Beyond possible phagocytic interactions, participants of the endocytic pathway were identified, such as the adaptor protein-2 (AP-2) complex. ARAP1 has been implicated in the past to regulate the endocytosis of EGFR and interact with another clathrin-associated member of the AP family, AP-3.^4,22^ Consistent with this possibility, many elements of clathrin (CLTA, CLTB, CLTC) and clathrin adaptors were identified (CLINT1, EPN1, HIP1, STON2).^32^,^33^ Unconventional myosin 6 (MYO6) and many of its formerly validated interactants were precipitated. These interactants include cargo adaptors DAB2 and TOM1 and the entirety of the DOCK7-Induced Septin disPlacement (DISP) complex (MYO6, DOCK7, LRCH3). These results are summarized in Table 1.

**Table 1.**
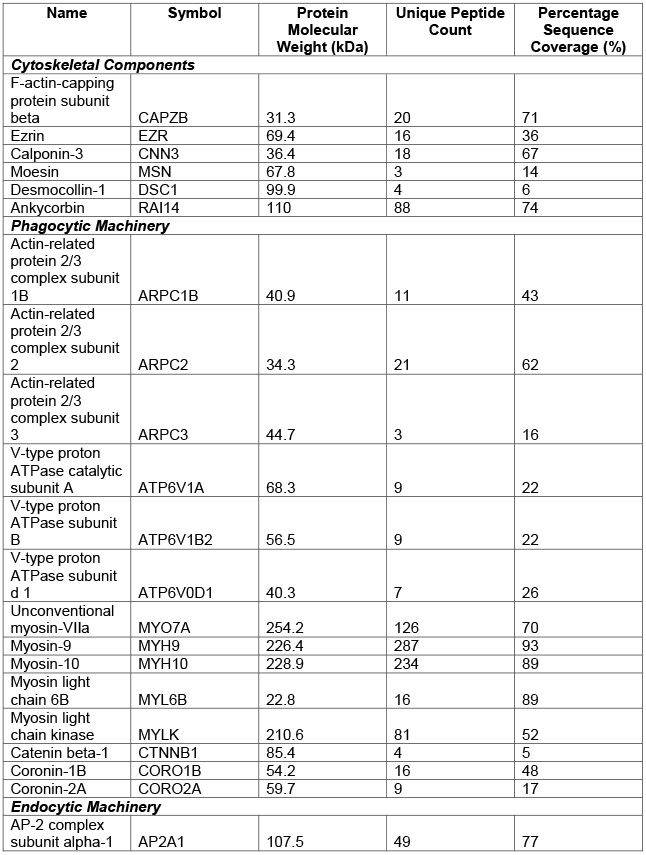
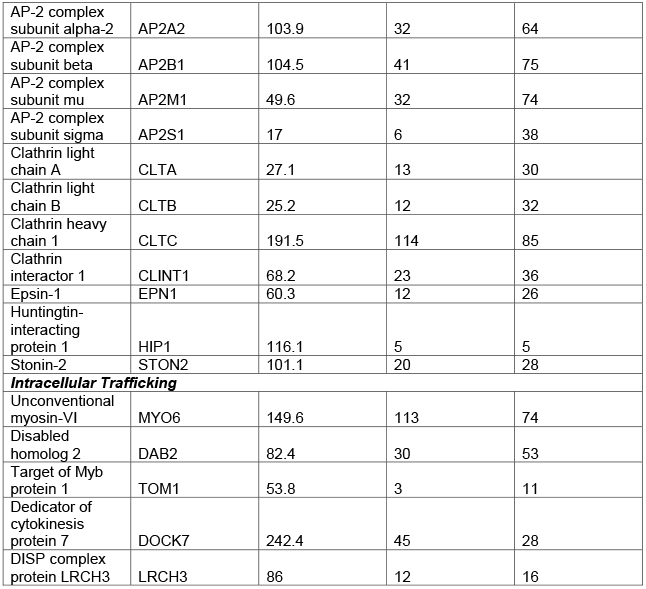
Putative ARAP1 binding partners. Proteins identified by LC-MS/MS analysis are shown above, segregated by cellular function. Protein name, symbol, and molecular weight are provided, as well as number of unique peptides and percentage of sequence coverage detected by mass spectrometry.

Several established Arap1 binding partners were immunoprecipitated but just missed criteria for inclusion after stratification. SH3 domain-containing kinase-binding protein 1 (CIN85) was detected in 2 of 5 experimental samples and in none of the controls. CIN85 has been formerly validated as an ARAP1 interactor by the yeast two-hybrid system and LC-MS/MS analysis.^22,34,35^ Cell division control protein 42 (Cdc42) was detected in 3 of 5 experimental samples and was significantly enriched compare to control IP. However, only 1 unique peptide was identified, preventing its inclusion. Ras-related C3 botulinum toxin substrate 1 (Rac1) was also identified in 3 of 5 experimental samples but lacked the enrichment required for its inclusion. Both Cdc42 and Rac1 were near absent in control samples. Cdc42 and Rac1 are key drivers of actin polymerization in RPE phagocytosis.^36^ Notable proteins failing inclusion criteria are summarized in supplementary table 1.

### *Arap1^-/-^* Mice Demonstrate RPE Phagocytic Defect

As LC-MS/MS analysis revealed a significant number of interactants related to phagocytic machinery, we evaluated *Arap1*’s role in OS phagocytosis. Retinal sections of *Arap1^-/-^* and *Arap1^+/+^* littermates were stained with fluorescent antibodies against rhodopsin (green) for rod quantification and M- and L- opsins (red) for cone quantification with DAPI counterstaining (blue) to better visualize the RPE layer. To account for circadian variation in RPE phagocytosis, mouse sacrifice was standardized to 1.5 hours after light onset.^37^ At postnatal day 24, *Arap1^-/-^* mice demonstrated a reduction in rod phagosomes (18.4 ± 2.7 per 200 μm retina) compared to their WT littermates (79.1 ± 13.3 per 200 μm retina) (Fig. 7A,7B,7E). Cone phagosomes, however, were comparable between the two groups (*Arap1^-/-^*:13.2 ± 3.7 per 2 mm retina, Arap1^+/+^: 10.5 ± 1.7 per 2 mm retina) (data not shown).

**Figure 7.**
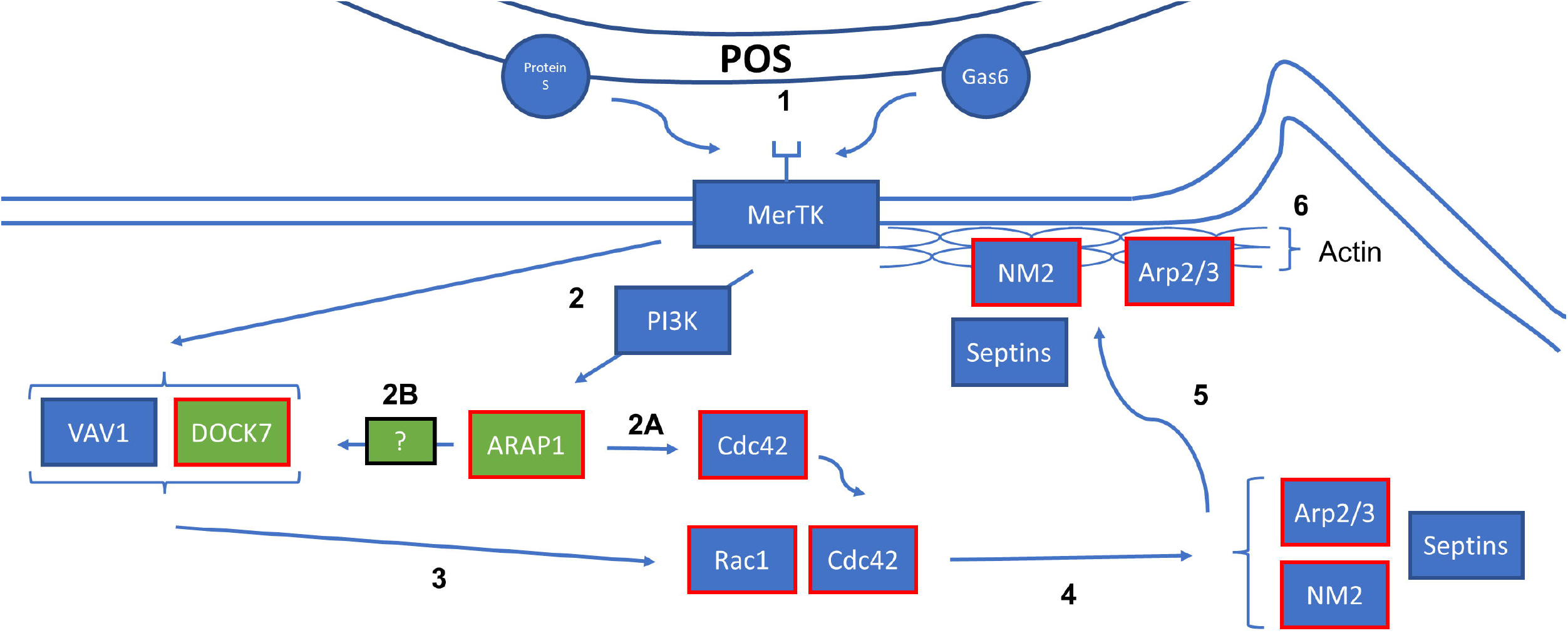
Reduction of RPE rod phagocytosis in Arap1^-/-^ retinas. Immunohistochemistry using anti-rhodopsin (green) and anti-M and anti-L opsin (red) was performed to quantify rod and cone RPE phagosomes, respectively, with DAPI counterstaining (blue) to visualize the RPE nuclei. Only merged images are shown. Rod phagosomes (white arrows) were counted in eyes sectioned at postnatal day 24 in both *Arap1^-/-^* mice and wild type litter mates (**A,B**). Scale bar (black) represents 20 um. *Arap1^-/-^* retinas demonstrated reduced numbers of rod phagosomes (18.4 ± 2.7 per 200 um retina) compared to wild type littermates (79.1 ± 13.3 per 200 um retina) (**E**). Phagosomes were also quantified in postnatal day 16 *Arap1^-/-^* and wild type littermates (**C,D**). *Arap1^-/-^* retinas at postnatal day 16 also demonstrated reduced numbers of rod phagosomes (18.2 ± 2.8 per 200 um retina) compared to their wild type littermates (73.1 ± 5.8 per 200 um retina)(**F**). Cone phagosomes (arrow in **D**) were comparable between *Arap1^-/-^* and wild type retinas at both postnatal day 24 and 16 (data not shown). Degenerative changes of *Arap1^-/-^* retinas are shown in (**G**). Vacuoles (arrowheads) and OS debris whorls (arrows) are frequently observed in the OS layer. These changes are absent in wild type retinas. (n = 3 each group, *P < 0.05, error bars represent SE)

However, *Arap1^-/-^* retinas already experience significant photoreceptor degradation by postnatal day 24. Compared to *Arap1^+/+^* retinas, the *Arap1^-/-^* rod OS were shorter and experienced extensive vacuolization. Whorls of membrane debris were also observed within the RPE-OS interface in mutant retinas (Fig. 7G). As such, OS phagocytosis of pups at eye-opening (postnatal day 16) were also quantified to confirm that loss of phagosomes was due to an intrinsic phagocytic defect. Analysis of postnatal day 16 retinas revealed markedly less degeneration, though occasional vacuolization was still observed. Measurement of postnatal day 16 RPE phagocytosis revealed a similar reduction in rod phagosomes in knockout retinas compared to *Arap1^+/+^* littermate retinas (*Arap1^-/-^*:18.2 ± 2.8 per 200 μm retina, *Arap1^+/+^*: 73.1 ± 5.8 per 200 μm retina) (Fig. 7C,7D,7F).

## DISCUSSION

The findings of our study reveal that *Arap1* expression in RPE is essential for photoreceptor survival due to its function as a critical interactant of RPE phagocytic machinery. *Cralbp-Cre Arap^tm1c/tm1c^* and *Vmd2-Cre Arap1^tm1c/tm1c^* mice recapitulate the phenotype originally observed in *Arap1^-/-^* mice as confirmed by histopathology, OCT, and fundus analysis. The milder degeneration observed in the *Cralbp-Cre Arap^tm1c/tm1c^* mice is explained by the relative reduction in RPE *Cre* function in this line, possibly secondary to reduced Cre expression by the *Cralbp* promoter in RPE cells (Fig. 2E). As suggested by phagocytic machinery identified in LC-MS/MS analysis of ARAP1 co-immunoprecipitation and confirmed with immunohistochemistry, the mechanism to these degenerative changes is the significant reduction in RPE rod OS phagocytosis secondary to *Arap1* loss. Rod phagosomes were generated by *Arap1^-/-^* RPE at approximately 25% of the rate of *Arap1^+/+^* RPE. This failure was accompanied by significant dysmorphic changes in the rod OS such as extensive vacuolization and whorls of OS membrane observed in the OS-RPE junction. It is somewhat unexpected that only rod phagocytosis was impeded while cone phagocytosis was comparable to controls. However, these abnormalities in phagocytosis do mirror the functional impediments we saw on ERG analysis of *Arap1^-/-^* mice previously, such that scotopic impairment appeared sooner and with greater severity than photopic impairment.^3^ Furthermore, previous studies suggest phagocytosis of rod and cone outer segments by RPE may be differentially regulated.^14^

Though *Arap1* is expressed in Müller glia, its function does not seem to be essential for maintaining photoreceptors, as corroborated by analysis of *Glast-Cre Arap1^tm1c/tm1c^* mice. Müller glia do however require *Arap1* for structural function. Ultrastructural analysis of *Glast-Cre Arap1^tm1c/tm1c^, Cralbp-Cre Arap1^tm1c/tm1c^*, and *Arap1^-/-^* retinas reveal discontinuity of the ELM absent in wild type and -*Cre Arap1^tm1c/+^* retinas. The ELM is a network-like series of junctional complexes between Müller glia and photoreceptors essential for maintaining structural integrity of the retina.^38^ Adherens junctions (AJ) are composed of E-cadherin, p120-catenin, β-catenin and α-catenin.^39^ As reviewed by Hartsock et al., p120-catenin has been shown to interact with Rho family GTPases to regulate actin cytoskeletal dynamics.^39^ Given its ability to regulate Rho GTPases, ARAP1 may regulate AJ-cytoskeleton communication in some capacity.

LC-MS/MS analysis of anti-ARAP1 immunoprecipitate has revealed several avenues of interest in uncovering both the specific function of ARAP1 in RPE cells and mechanisms of RPE phagocytosis. Considering the diversity of ARAP1’s functional domains, it is not surprising that several different processes are implicated by the results of mass spectrometry analysis. Given that interruption of photopigment (11-cis retinaldehyde) recycling can cause retinitis pigmentosa, it is noteworthy that no proteins related to this pathway were identified.^9^

Many of the proteins identified were involved in the clathrin-mediated endocytic pathway such as the AP-2 complex, clathrin elements, endocytic adaptors, and MYO6 and its interactants.^40,41^ DAB2’s inclusion is particularly interesting as it has been shown to bind both the AP-2 complex and MYO6, bridging endocytosis and actin cytoskeletal regulation - two processes in which ARAP1 has been implicated.^42^ DAB2 has also been shown to bind CIN85, an established ARAP1 interactant, to target CIN85 to clathrin-coat assembly for EGFR endocytosis.^43^ CIN85, AP-1, −2, and −3 complexes, clathrin, and ARAP1 have been shown to colocalize to during EGFR endocytosis.^35^ Furthermore, CIN85/ARAP1 likely affect targeting of the pre-early endosome, the endocytic stage at which DAB2 and MYO6 are active.^35,44^ Though unable to meet inclusion criteria, CIN85 is a notable protein identified in our LC-MS/MS analysis (Supp. Table 1). Given that MYO6/DAB2 and ARAP1/CIN85 interactions all function in the pre-early endosome and the interactions between DAB2/CIN85, their identification in our co-immunoprecipitation may suggest that these proteins function under one unified mechanism. As the mechanisms targeting MYO6 to DAB2 in the nascent endosome remain unelucidated, it is tempting to posit that ARAP1/CIN85 may function to fulfill this role. Ultimately, these results echo past conclusions of ARAP1’s involvement in clathrin-mediated endocytosis, specifically in the pre-early endosome.

LC-MS/MS revealed significant interactants of RPE phagocytosis, which is mechanistically similar to the Fcγ receptor (FcγR) phagocytosis system as reviewed by Kevany et al.^36^ Accordingly, RPE phagocytosis is comprised of several sequential steps: recognition, binding, internalization, intracellular trafficking, and digestion. Components of RPE phagocytic machinery have been identified previously. Mer tyrosine kinase (MerTK) and its ligands Gas6 and Protein S are involved in the internalization of shed OS membranes, adhesion receptor αvβ5 integrin regulates recognition/binding, and scavenger receptor CD36 facilitates internalization.^36^

Beyond receptor interactions, several cytoplasmic proteins have also been linked to RPE phagocytosis. Knockout of myosin VII and annexin A2 leaves engulfed phagosomes localized in the apical region of RPE cells while WT RPE traffic phagosomes to the basal region.^36,45^ Even further downstream in the phagocytic pathway, knockout of melanoregulin, a putative membrane fusion regulator protein, demonstrates a normal number of phagosomes generated but without normal decline, suggesting impairment in phagosome digestion post-engulfment.^36^

Looking at how failures at each step affect phagosome analysis, it appears that *Arap1* loss most likely causes a failure at either binding or engulfment. Were there a failure in intracellular trafficking, phagosome counts would be normal, though with abnormal localization. Similarly, a failure in digestion would generate normal phagosome counts that fail to reduce appropriately. Supporting this theory is the Royal College of Surgeons (RCS) rat, a model for retinitis pigmentosa. Similar to *Arap1^-/-^* mice, RCS rats also develop progressive photoreceptor degeneration beginning with rods, despite proper photoreceptor development, secondary to RPE phagocytic defect.^46^ The mutant locus (*rdy*) of the RCS rat was identified to correspond with the gene *Mertk* and reconstitution of the gene rescues the RCS phenotype.^47,48^

Reinforcing the parallel, *Mer* knockout mice share striking similarities to *Arap1^-/-^* mice. Histopathology of *Mer* knockout mice reveal progressive loss of the ONL, characterized by early rod loss, and vacuole formation in the OS.^49^ ERG analysis revealed a progressive impairment of both scotopic and photopic responses, with scotopic impairment appearing earlier and at greater severity.^49^ Most importantly, they demonstrate a marked reduction in RPE phagosomes compared to WT counterparts. Given the apparent similarities between the RCS rat, *Mer* knockout mice, and *Arap1^-/-^* mice, *Arap1* loss would seem to disrupt the phagocytic process through a similar mechanism as *Mertk* loss.

Mass spectrometry analysis corroborates this possibility. Many proteins identified have clear roles in the cytoskeletal dynamics of RPE phagocytosis. The ARP2/3 complex and beta-catenin play essential roles in actin polymerization and organization during phagocytosis.^36^ Even more fascinating are the components and interactants of nonmuscle myosin ii (NM2) identified: MYH9, MYH10, MYL6B, and MYLK.^50–53^ NM2 has been shown to be recruited downstream of MerTK signaling and is essential for RPE engulfment of POS. Following that observation, it is hypothesized that NM2 and MerTK may be members of a protein complex that governs RPE phagocytic engulfment.^30^ Interestingly, the motor proteins of isoforms NM2A and NM2B (MYH9 and MYH10, respectively) were both identified. Though depletion of either isoform impede RPE phagocytosis, NM2A has been identified as a specific interactant of MerTK per immunoprecipitation and mass spectrometry analysis.^30^ This result is in line with the understanding that NM2 mediates cellular protrusion, a process essential for the formation of phagocytic pseudopodia.^54^ Even more convincing that NM2 is likely involved in the phenotype of *Arap1* loss is NM2’s role in stabilizing AJ integrity.^55,56^ Recalling the ELM abnormalities seen in *Arap1^-/-^* and *Glast* and *Cralbp-Cre Arap^tm1c/tm1c^* mice, involvement of NM2 would not only explain the phagocytic defect we see in RPE cells, but also the abnormal AJ seen in Müller glia. Similar to their dynamic in RPE phagocytosis, NM2A and NM2B play complementary roles in AJ formation and maintenance.^56^ Though interaction between ARAP1 and NM2 has yet to be documented, another ArfGAP family member, ASAP1, has been previously shown to bind NM2A.^26^

Cdc42 and Rac1, two proteins that were identified but unable to meet inclusion criteria, have both been shown to be upstream of NM2 recruitment.^57,58^ Furthermore, they have been shown to activate under mechanical stress in E-Cadherin expressing cells, a NM2-regulated process.^57,59^ The identity of the guanine exchange factor (GEF) regulating Cdc42 and Rac1 phagocytic signaling has been a subject of debate.^36^ The answer may lie in the DISP complex that was identified in our LC-MS/MS analysis.

The DISP complex (LRCH3, MYO6, DOCK7) has been shown to regulate the organization of septins, a family of filament-forming, GTP-binding proteins essential for phagosome formation in FcγR phagocytosis.^40,60^ Furthermore, members of the septin family are essential in scaffolding for the formation of NM2 fibers.^61^ Notably, DOCK7 has been shown to be a GEF for both Rac1 and Cdc42.^62–64^. Though Vav1, a downstream effector of MerTK, has been shown exhibit GEF activity for Rac1, its activation of Cdc42 has been debated.^65,66^ Furthermore, Cdc42 has been shown to be essential for septin ring formation.^67^ Given DOCK7’s GEF activity against Cdc42, Cdc42 could be the downstream effector of the DISP complex in septin arrangement and ultimately NM2 recruitment. If this were the case, then the DISP complex would provide an answer for the source of both GEF activity against Cdc42 and septin regulation during phagocytosis.

ARAP1’s role in endocytosis has been explored in the past, particularly in the role of EGFR recycling.^4^ Given identification of both the AP-2 complex and MYO6 interactome, ARAP1 appears to fulfill this role in RPE. Furthermore, ARAP1’s GAP activity against Arf and Rho family G proteins has established a clear role for ARAP1 in cytoskeletal dynamics - consistent with the many elements of the RPE cytoskeleton we identified.^5^ However, this is the first instance establishing a role for ARAP1 in phagocytosis, as corroborated by the RPE phagocytosis defect observed in *Arap1^-/-^* mice and the numerous elements of RPE phagocytic machinery identified. The reduced number of phagosomes seen in *Arap1^-/-^* retinas as well as the phenotype observed in *Arap1*-deficient mice seem to indicate a likely failure in phagosome engulfment, as corroborated by the RCS rat and *Mer* KO mice. Supporting this theory are the numerous elements of NM2 and the DISP complex that were identified, as well as the ARP2/3 complex which has been well-described in its role in engulfment and cellular protrusion.^54,68^

Though Rac1 and Cdc42 did not meet criteria for inclusion as putative interactants, suspicion remains high for potential interaction with ARAP1 - particularly Cdc42. Compellingly, ARAP1’s effects on filopodia formation and F-actin organization have been shown to be mediated through its ArfGAP activity against Cdc42.^5^ It is understood that Cdc42 and Rac1 play independent, though spatiotemporally coordinated roles in FcγR phagocytosis.^54^ Given the essential regulation of Cdc42 during NM2-mediated protrusion, it is tempting to posit ARAP1 participates in this process.^54^ Indeed, there is a possibility ARAP1 not only participates, but may do so via DOCK7. DOCK7 regulation is largely unknown, however there is circumstantial evidence that it may involve phosphatidylinositol (3,4,5) trisphosphate (PIP3). DOCK7 has been shown to be an essential regulator of neuronal polarity and axon growth, a PI3K/PIP3 dependent process.^64,69,70^ Given ARAP1’s PIP3-depedent ArfGAP activity, it is possible that PI3K/PIP3 effects on DOCK7 are mediated through ARAP1.^5^ Consistent with this possibility, MerTK has been shown to activate PI3K and PI3K has been shown to be essential for large particle phagocytosis.^71,72^ An additional possibility is that ARAP1 may exhibit RhoGAP activity to inactivate Cdc42/Rac1. RhoGAPs have been noted to accumulate in a PI3K-dependent fashion to large phagocytic cups and Cdc42/Rac1 inactivation is essential for formation of these large phagosomes.^72^ A proposed model for these interactions is shown in Figure 8. Of course, future investigations will be necessary to confirm these suspicions. However, these results provide a novel role for ARAP1 as an essential component of RPE phagocytosis and further elucidate the mechanisms driving this complex process.

**Figure 8.**
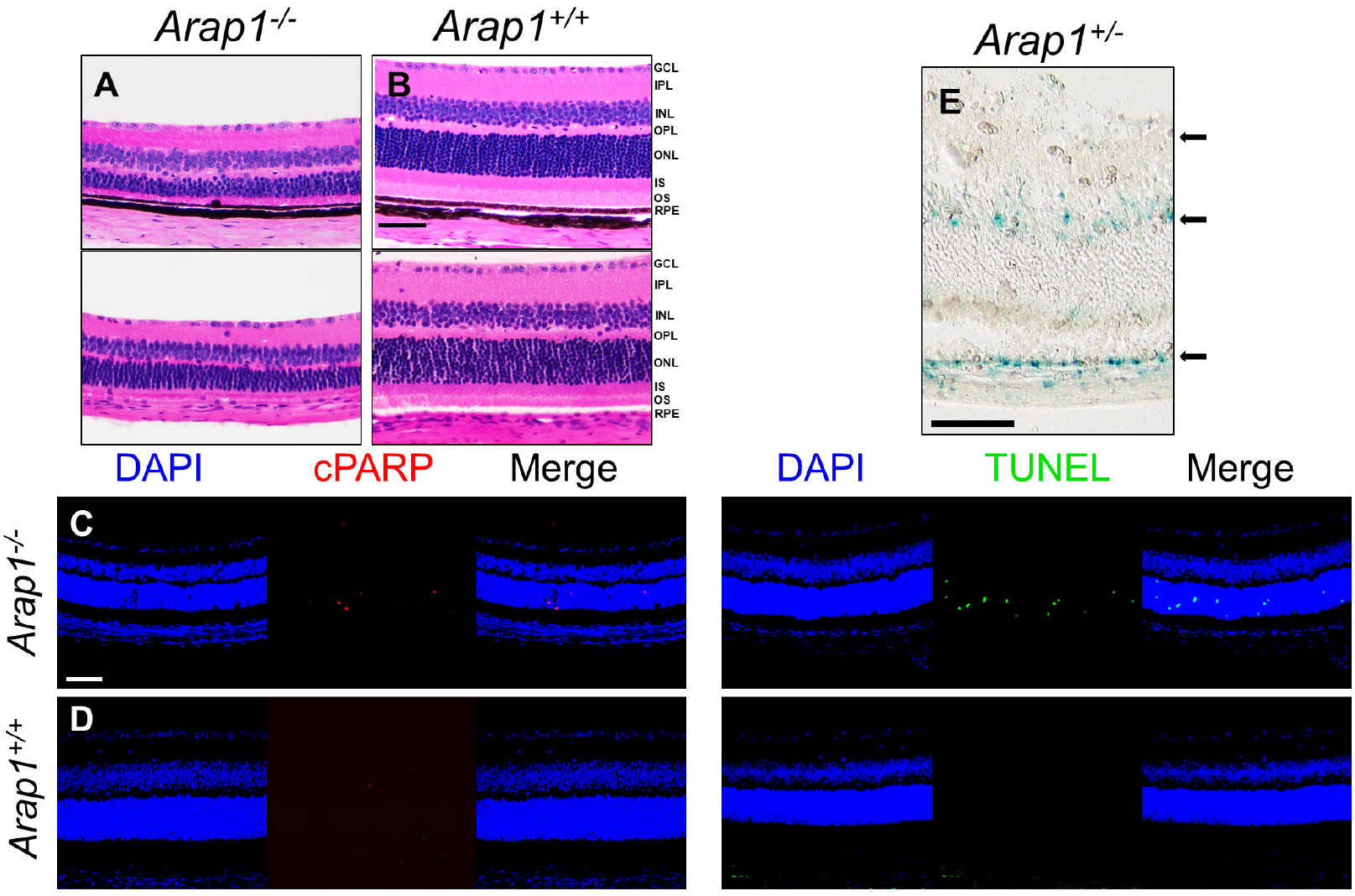
Proposed Model of RPE Phagocytosis. A visual summary of interactions reviewed in the discussion is shown. Proteins established in the MerTK phagocytosis cascade are filled blue, while proposed interactants are filled green. Red outlines dictate that the protein, or components of the protein, were identified by mass spectrometry analysis. 1. POS bound by MerTK ligands Protein S and Gas6 bind MerTK receptor 2. MerTK-ligand binding recruits and activates GAP (e.g. ARAP1) and GEF (e.g. DOCK7, VAV1) activity proteins. 2A. ARAP1 localizes Cdc42 or 2B. ARAP1 activates DOCK7 by endogenous binding or an unknown intermediary via its ArfGAP domain 3. GEF activity proteins activate G protein Rac1 and Cdc42 4,5. Rac1 and Cdc42 recruit NM2, septins, and drivers of actin polymerization (e.g. the Arp2/3 complex) from the cytoplasm and localize it to the site of MerTK. 6. Localization of NM2 with cortical actin provides structural reinforcement for actin remodeling (e.g. via Arp2/3) and subsequent phagocytosis.

---

**Supplementary table 1.** Notable proteins from LC-MS/MS analysis. The proteins above are proteins with notable interactions with ARAP1 corroborated by literature search but were unable to meet inclusion criteria. Protein name, symbol, molecular weight are provided, as well as number of unique peptides and percentage of sequence coverage detected by mass spectrometry.

| Name | Symbol | Protein Molecular Weight (kDa) | Unique Peptide Count | Percentage Sequence Coverage (%) |
| --- | --- | --- | --- | --- |
| SH3 domain-containing kinase-binding protein 1 | CIN85 | 73.1 | 1 | 2 |
| Cell division control protein 42 | CDC42 | 21.2 | 1 | 11 |
| Ras-related C3 botulinum toxin | RAC1 | 21.4 | 2 | 18 |
| substrate 1 |  |  |  |  |

Supplemental Figure 1 – Characterization of Cre^tm1c/+^ mice. *Cre^tm1c/+^* mice were analyzed with fundus photography, histology, and OCT analysis at 3 months of age (Glast-Cre, Cralbp-Cre) and 1 month of age (Vmd2-Cre). Fundus photography, histopathology, and OCT analysis were all unremarkable. Quantification of retinal layers is shown in Figure 3C, 3G, and 3K.

Supplemental Figure 2. Uncropped Blots shown in Figure 7. Immunoblot of WT and Arap1^-/-^ mice in biological triplicate is shown (A). Immunoblot of FRPE lysate, anti-ARAP1 immunoprecipitate, and goat IgG immunoprecipitate is shown (B).

## Acknowledgements

Special thanks to Dr. Henry Ho Ph.D., Monica Motta, Brad Shibata and the NEI Core Microscopy Lab at UC Davis, and Dr. Gabriela Grigorean Ph.D. and the UC Davis Proteomics Center.

Research generously supported by mentored career development grant NIH NEI K08 EY027463

The authors declare no competing interests.

