## Supplementary figures and images for "*Arap1* Loss Causes RPE Phagocytic Dysfunction and Subsequent Photoreceptor Death"

### Supplemental Figure 1

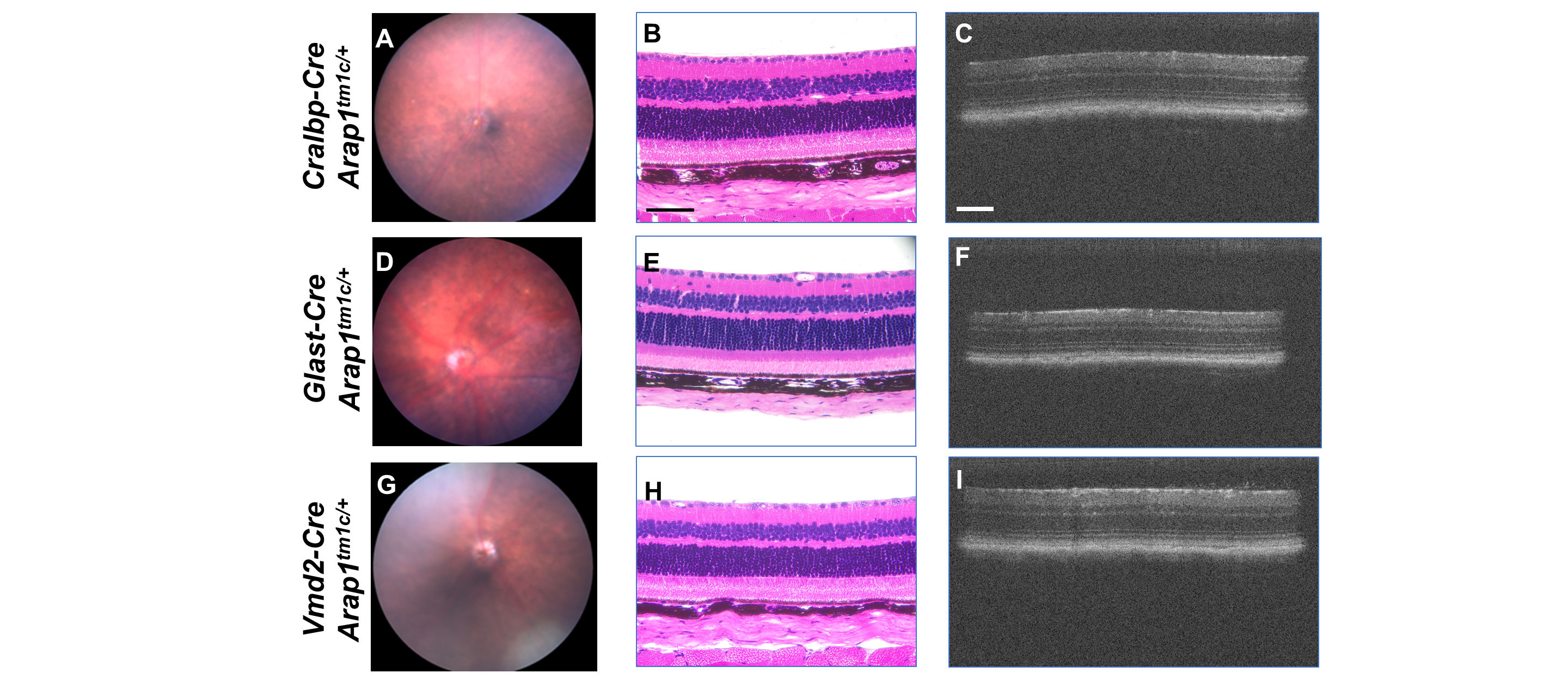

### Supplemental Figure 2

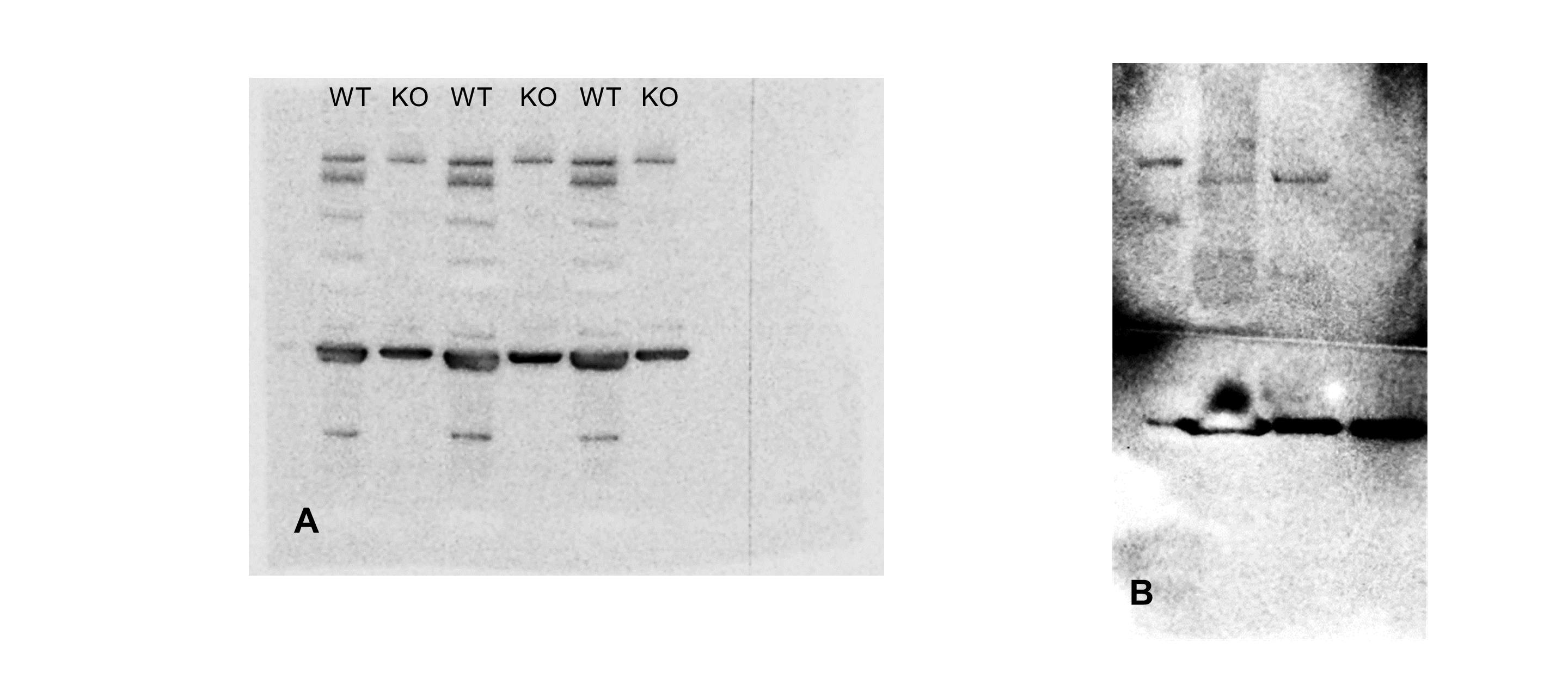
